# Exome sequences of toque macaques (*Macaca sinica*) of Sri Lanka reveal many amino acid changes

**DOI:** 10.1101/740761

**Authors:** Nilmini Hettiarachchi, Naoki Osada, Hirofumi Nakaoka, Takashi Hayakawa, Ituro Inoue, Naruya Saitou

## Abstract

Macaques are one of the most widely used model organisms in biomedical research. *Macaca mulatta* and *M. fascicularis* are currently being used for toxicology, HIV, diabetes, neuroscience and psychiatric and psychological disorder researches. Many studies have been conducted on *M. mulatta* and *M. fascicularis* genomes for this purpose in order to understand the genomic properties of these species. Several *M. fascicularis* individuals from different geographical locations and also *M. mulatta* genomes have been sequenced and studied in depth for the purpose of understanding the phylogenetic and evolutionary relationships between various macaque populations. But still a gap in knowledge remains for other macaque populations such as the *Sinica* group. In this study for the very first time, we sequenced the exome of toque macaques (*M. sinica*), an endemic island population of Sri Lanka. Here we confirmed that *M. sinica* and *M. thibetana* cluster together and are closely related, also the three distinct phylogenetic groupings of *fascicularis* and *sinica*. We also found that *M. sinica* has less number of polymorphisms with respect to the reference genome *M. mulatta* signifying the smaller and restricted population size of this species.

## 1. Introduction

Macaques are Old World monkeys that belong to subfamily Cercopithecoidea and tribe Papionini. Humans and macaques shared a common ancestor about 25 million years ago (Kumar and Hedges 1998). They are predominantly distributed in southern and eastern parts of Asia, except Barbary macaques which distribute in northern parts of Africa, specifically regions of Algeria and Morocco (Lavieren 2012). Macaques have been introduced to some countries in the world where they did not exist. One such example is Mauritius Island, where *fasciclaris* macaques were introduced by sailors from Java in the 17^th^ century (Sussman and Tattersall 1981). Initially macaques were classified into four distinct groups by Fooden (1976), namely *silenus*-*sylvanus, fascicularis, arctoides*, and *sinica*. Over the years there has been new developments regarding the categorization of the macaques. Currently the macaques are grouped in to three main categories, namely *fascicularis, sinica* and *silenus* (Delson 1980; Hayasaka et al. 1996). The *fascicularis* group consists of four species following Fooden (2006); *M. fascicularis, M. mulatta, M. cyclopis*, and *M. fuscata.* The *sinica* group consists of *M. sinica, M. radiata, M. assamensis*, and *M. thibetana* (Hayasaka et al. 1996). The *silenus* group contains lion tailed macaques (*M. silenus*), pig tailed macaques (*M. nemestrina*) and stump tailed macaques (*M. arctoides*) (Hayasaka et al. 1996). Li et al. (2010) proposed four groups of macaque species; *sylvanus, silenus, sinica* and *fasciculari*s based on 358 Alu insertion polymorphisms.

Macaques are one of the mainly used non-human primates in biomedical research, in toxicology, research in HIV, diabetes, neuroscience and psychiatric and psychological disorders. This is because macaques show high genetic similarity to humans (94%; Osada et al. 2015). Rhesus macaque (*M. mulatta*) is the most widely used for research purposes, while crab-eating macaques (*M. fascicularis*), pig-tailed macaques (*M. nemestrina*) and Japanese macaques (*M. fuscata*) are also used. *M. fascicularis* of Mauritius Island are commonly used for research purpose, as they are genetically highly heterogeneous which make them valuable in testing responses to various drugs (Osada et al. 2010; Yan et al. 2011; Osada et al. 2015). Fan et al. 2014; Osada et al. 2015; Fan et al. 2018 also found that within individuals the proportion of heterozygous nonsynonymous polymorphisms are higher compared to synonymous polymorphisms in Mauritius *M. fascicularis* due to strong population bottle neck.

The main interest of this study is *M. sinica*, commonly known as toque macaques for the tuft of hair on top of the head which resembles a hat. This species belongs to the *sinica* group and is endemic to Sri Lanka. There are three different subspecies within the island based on their geographical distribution. *M. sinica sinica, M. sinica aurifrons*, and *M. sinica opisthomelas* are found in dry zone, wet zone and highlands of Sri Lanka, respectively. These macaques have been studied since 1960s with regards to hematology (Hawkey et al. 1978; Jain 1986), pathogens, parasites and viruses (Dissanaike 1965; Dewit et al 1991). Hoelzer et al. (1994) identified highly diverged mitochondrial DNA haplotypes in *M. sinica.* However, *M. sinica* genome has not yet been studied from a comparative genomics perspective. The main reason for this is the unavailability of genome sequences of *M. sinica*. Here we attempted to fill the gap of lacking knowledge and also data by sequencing the exome of *M. sinica* in order to expand the understanding of toque macaque genome from an evolutionary and comparative genomics point of view.

## 2. Materials and Methods

### 2.1. DNA sequencing

We extracted the genomic DNA from muscle tissue of a male *M. sinica* reared at Japan Monkey Centre, Inuyama, Japan (specimen number: Pr6182) using DNeasy Blood & Tissue Kit This individual is an offspring of first-degree inbreeding (father and daughter) and the father and the daughter’s mother are also captive-born but their origin cannot be traced more. The library construction and sequencing was conducted at Human Genetics Laboratory at National Institute of Genetics, Mishima, Japan. Three µg of DNA was sheared into fragments with length of 150-200 bp using Covaris S2 focused-ultrasonicator. Sequencing library was constructed with SureSelect XT reagent kits. Target enrichment was constructed with SureSelect Human All Exon V5 + lncRNA kit. The library was sequenced with Illumina HiSeq 2500 platform, in rapid run mode with 2 × 150-bp paired-end module. The quality check of the sequence reads was performed by using FastQC (Andrew 2010) and all reads were above Q30 quality level.

### 2.2. Other macaque genomes used in the comparative analysis

We used eight macaque genomes to compare with toque macaque exome data; three *M. mulatta* (SRR1927155, SRR1929293, SRR1950974) (Xue et al. 2016), one Vietnamese *M. fascicularis* (SRA023855) (Yan et al. 2011), one Malaysian *M. fascicularis* (Higashino et al. 2012), two Mauritian *M. fascicularis* (SRR1564753, SRR1564755) (Ericsen et al. 2014), and one *M. thibetana* (SRR1024051) (Fan et al. 2014). These 8 individuals (section 2.3) were used separately for variant calling to determine effects of SNPs on genes of all 7 individuals except reference. The reason for this separate variant calling (stage-1) with rhesus macaque as reference is the availability of better quality annotation data for genes for rhesus macaque that would help in determining the effects of variants on these genomes.

In order to obtain an abstract view on genetic relationships between species, we again called variants (stage-2) using *M. nemestrina* as the reference (section 2.4) with an addition of several other newly sequenced species (*M. assemensis* and *M. arctoides*). This was to determine phylogenetic relationships between these individuals via phylogenetic tree and Principal Component Analysis (PCA). But these species were not considered for down-stream analyses on SNP effect.

### 2.3. Stage-1 variant calling for 8 macaque individuals with *M. mulatta* as reference

Variant calling was performed simultaneously on *M. sinica, M. thibetana*, two *M. fascicularis* (Mauritius), *M. fascicularis* (Malaysian), *M. fascicularis* (Vietnam), *M. mulatta* (Chinese), two *M. mulatta* (Indian) genomes. Genotypes were called with the Best Practice pipeline of Genome Analysis Tool Kit (GATK) (DePristo et al. 2011).

### 2.4. Phylogenetic relationship between the species used in the study with stage-2 variant calling with *M. nemestrina* as reference

In order to construct an elaborate phylogenetic tree with many recently sequenced macaque species, we remapped the exome reads from *M. sinica* to the *M. nemestrina* (GCA_000956065.1) genome, which would work as an out-group. The choice of reference genome here also removes mapping bias that might occur by using the rhesus macaque genome as the reference, because *M. nemestrina* is expected to be equally distant from the *sinca* and *fascicularis* groups. In addition, we added two recently-determined genome sequences from the *sinica* group: one *M. assemensis* (SRR2981114), and one *M. arctoides* (SRR2981139) (Fan et al. 2018) into the analysis. The data of Malaysian *M. fascicularis* was removed from the phylogenetic analysis because the reads of the samples were determined by a SOLiD sequencer and would not be suitable for mapping to a new reference genome. The Neighbor joining tree (Saitou and Nei. 1987) was constructed with distance matrix determined based on genotype data for the 11 individuals used for this analysis, using the Phylo package in R.

### 2.5. Principal Component Analysis (PCA)

Stage-2 variant calling variants that were found in all 10 individuals except the reference (*M. nemestrina*) (section 2.6) were used to construct PCA. The filtering and processing of data for making ped and map files was performed by custom made perl scripts. PCA was constructed with smartpca program in EIGENSOFT software package (Patterson et al. 2006).

### 2.6. Divergence time estimation for sinica and fascicularis lineages

Genotype data of all variant and invariant sites were used to calculate genetic distances between individuals. Evolutionary rate of 1×10^−8^ per site/per generation and generation time of 11 years (Xue et al. 2016) was assumed for the divergence time estimation.

### 2.7. Effect of SNPs

We annotated all variants and also the probable effect or in other words functional consequences of variants by SnpEff (Cingolani et al. 2012). Gene annotations are based on Mmul_8.0.1.86. Further we classified the genes in which the high and low impact variants are located for all 9 individuals in the study. The high impact SNP genes were classified by Database for Annotation, Visualization and Integrated Discovery (DAVID) by Huang et al. (2009a, 2009b).

### 2.8 *M. Sinica* only heterozygous non-synonymous sites

We looked at the heterozygous non-synoymous sites found only in *M. sinica* genome. These sites were analyzed in detail for its effect on the genome. The possible functionality of the genes in which these sites are found and the disease conditions that can be caused due to mutations were further looked at.

## 3. Results

### 3.1. Exome sequencing of *M. sinica* and variant identification summary for the nine macaque individuals

The exome sequencing of the *M. sinica* was performed on Illumina HiSeq 2500 platform and more than 22,280,000 pair end reads were generated. The reads have been deposited in the National Center for Biotechnology Information (NCBI) Short Read Archive under accession number PRJNA546445. The average depth calculated for the mapped exome was 50x. A total of 394,202 variant sites were identified for the nine individuals in the study with rhesus macaque as the reference during stage-1 variant calling. *M. thibetana* and *M. sinica* had less number of polymorphisms. This less genetic diversity is expected in isolated island populations such as *M. sinica. M. sinica* is endemic to Sri Lanka which is an island in South Asia but the observation of less number of polymorphisms in *M. thibetana* requires further analyses which we will not be addressing during this study. The small restricted population size and isolation leads to inbreeding and then again to reduced genetic variation, as with the case of *M. sinica*. Also lower genetic diversity can lead to extinction of the species in the long run. With respect to the reference genome, *M. mulatta*, we identified 26,126 heterozygous sites (Supplementary Figure 1) which is relatively lower than for the other species in the analysis. *M. thibetana* shows a similar pattern to *M. sinica. M. thibetana* is also an endemic group to China (Fan et al. 2014). In general *M. sinica* and *M. thibetana* showed considerably high levels of homozygous sites (Supplementary Figure 1). Among other factors such as mutation rate, natural selection and patterns of recombination and linkage disequilibrium, excessive increase in homozygous sites can also be directly related to inbreeding (Gibson et al. 2006) due to isolation and small population size which can be observed in both *M. sinica* and *M. thibetana* populations. One important consideration is that the DNA sample we used for this study came from an individual that resulted from inbreeding (father-daughter inbred offspring). This causes the heterozygosity levels to drop to 75% in the offspring by considering inbreeding coefficient as 25%. In a case where 0.25 of inbreeding coefficient is considered our results show lower level of heterozygosity (Supplementary Table 1) than expected. Therefore in this case lower level of heterozygosity observed in the *M. sinica* individual should be due to reasons other than inbreeding.

We determined the heterozygous synonymous and nonsynoymous SNPs for each species or sample. Synonymous SNPs will result in same amino acid as the reference genome and if the SNP is nonsynonymous it will result in a different amino acid compared to the reference genome (Supplementary Figure 1). We also determined the ratio of nonsynonymous to synonymous variants (Table 1) for heterozygous sites. The ratio is slightly higher for *M. thibetana* and *M. sinica* compared to the *M. fascicularis* and *M. mulatta* individuals. This finding is plausible due to the smaller population size of *M. thibetana* and *M. sinica* macaques.

**Table 1.** Non-synonymous to synonymous ratios of the nine macaques. For ratio determination only the heterozygous sites were considered. The three groups in the study *Sinica, Mulatta* and *Fascicularis* are highlighted in colors blue, green and yellow respectively.

| Individual | Non-synonymous sites | Synonymous sites | Non-synonymous SNPs/Synonymous SNPs |
| --- | --- | --- | --- |
| *M. thibetana* (China) | 8105 | 8539 | 0.9491 |
| *M. sinica* (Sri Lanka) | 10315 | 11889 | 0.8676 |
| *M. mulatta* (China) | 16494 | 21693 | 0.7603 |
| *M. mulatta* 1 (India) | 14984 | 20439 | 0.7331 |
| *M. mulatta* 2 (India) | 15646 | 20991 | 0.7453 |
| *M. fascicularis* (Vietnam) | 19234 | 26498 | 0.7258 |
| *M. fascicularis* (Malaysia) | 18643 | 27275 | 0.6835 |
| *M. fascicularis* 1 (Mauritius) | 16643 | 21209 | 0.7847 |
| *M. fascicularis* 2 (Mauritius) | 17028 | 21539 | 0.7905 |

### 3.2. Genetic relations between populations with PCA

The macaque individuals showed clustering pattern where, *Fascicularis, Sinica* and *Mulatta* macaques were in three distinct groups (Supplementary Figure 2). Mauritian *Fascicularis* macaques were found to be genetically closer to the Vietnamese *Fascicularis* macaques. This result reconfirms findings by Osada et al. (2015). *M. thibetana M. sinica, M. arctoides, M. assemensis* clustered together in the PCA signifying *sinica* group.

### 3.3. Genetic relations between populations with phylogenetic tree

The phylogenetic tree that was constructed using 11 genomes which included newly sequenced stump-tailed macaque (*Macaca arctoides)*, Assamese macaque (*M. assamensis)* showed clustering of *M. sinica, M. thibetana* and *M. assamensi*s. This phylogeny also coincides with Fan et al. 2018 where stump-tailed macaque clusters close to *sinica* group. It is known that stump-tailed macaques have an unresolved genetic relationship with other macaque species, but our result here agrees with Fan et al. 2014 signifying that stump-tailed macaque is more closely related to *sinica* group than to *fascicularis* or *mulatta*. Residing with one of the primary objective of the study we confirm that Sri Lankan *M. sinica* and Tibetan macaques belong to the same phylogenetic group (Figure 1).

**Fig1.**
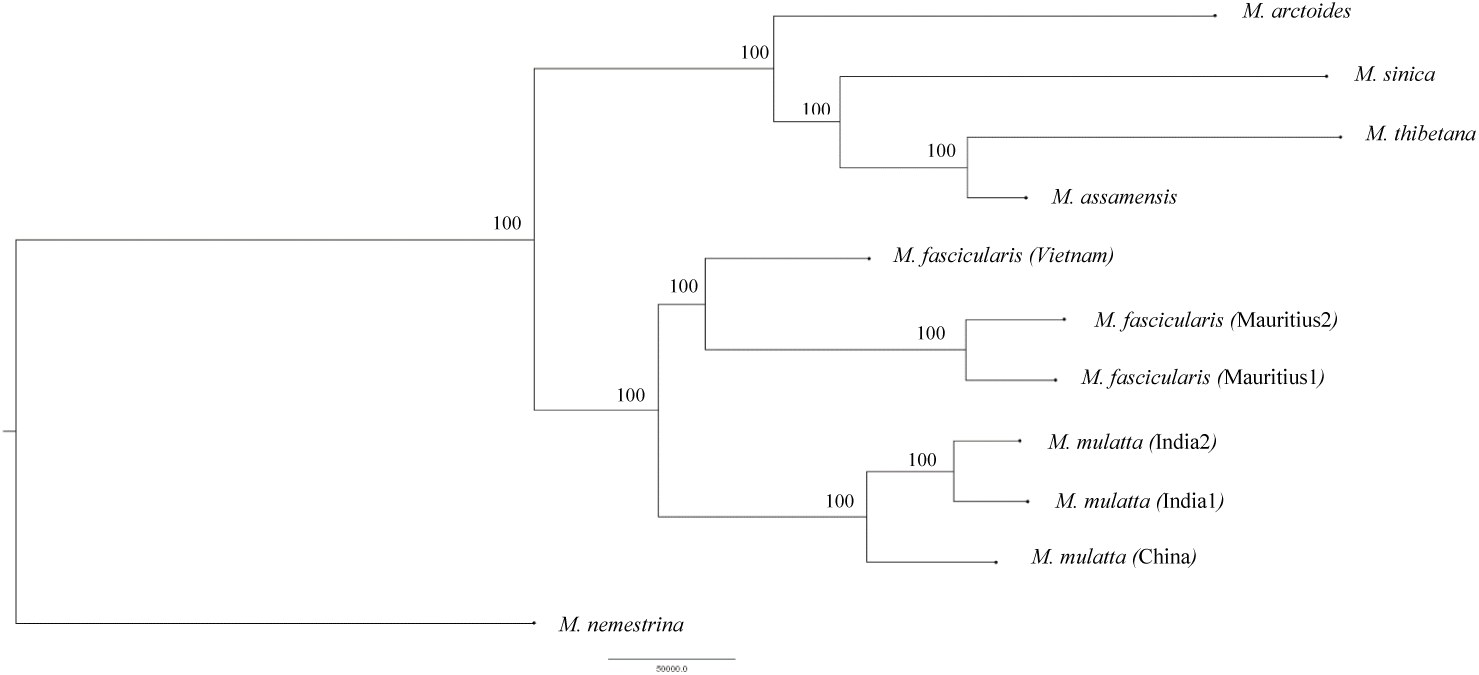
Phylogenetic relationships of the nine macaques. The tree was constructed based on the pair-wise distances calculated by genotype data

### 3.4. Divergence time estimation for *sinica* and *fascicularis* groups

*Fascicularis* and *Mulatta* individuals clustered together as the *fascicularis* group. *M. sinica* and *M. thibetana* individuals were clustered outside of the *Fascicularis* and *Mulatta* group (Figure 2). This finding is in congruence with Jiang et al. (2016) who reported that the *Fascicularis* group and the *Sinica* group clustered into two distinct groups based on 84 *Alu* insertion polymorphisms. Assuming a uniform evolutionary rate of 1×10^−8^ per site/per generation and a generation time of 11 years for macaques (Xue et al. 2016), the divergence time estimated for the *M. mulatta* and *M. fascicularis* group was approximately 1.3 million years which is close to the estimation of Osada et al. (2008), who estimated that *M. fascicularis* and *M. mulatta* shared a common ancestor about 0.9 million years ago. We also estimated that *sinica* and *fascicularis* groups diverged about 1.7 million years ago.

### 3.5. Effect of SNPs

We found that 2672 SNPs were annotated as HIGH impact by SnpEff4.3v across all 9 individuals in this study. The variant effects are classified into LOW and MODERATE effect by SnpEff4.3v. The majority of the variants belonged to MODERATE classification group. Interpretation of the LOW and MODERATE classifications is that they have lower phenotypic impacts on the gene. The numbers of variants in each group are provided in Table 2. We further classified the genes with high impact SNPs, and this analysis revealed several functional classification groups. Highest enrichment group contained cytochrome P450 (CYP) genes, second group contained genes with guanylate binding protein and septin. The third functional group contained MHC class I and class II genes. CYP enzymes are known to have an impact on drug metabolism of humans and macaques (Zuber et al. 2002). CYP class of enzymes is also involved in metabolism of certain anticancer drugs (Kivistö et al. 1995). Guanylate binding proteins are responsible for cell autonomous immunity and inflammasome activation (Man et al. 2016). The Major Histocompatibility region (MHC) plays a key role in immune response (Karl et al. 2013) and the third highest enrichment was related to this class of genes (Table 3).

**Table 2.** SNPs classified by SNPeff as high, low or moderate effect found in all 9 individuals.

| Classified Type | Number of sites |
| --- | --- |
| HIGH | 2672 |
| LOW | 179416 |
| MODERATE | 123178 |

**Table 3.** GO of genes for annotated high impact deleterious SNPs found in all 9 individuals. (A) Highest enrichment class of genes. (B) Second highest enrichment group of genes (C) Third highest enrichment group

| GO term | P-value |
| --- | --- |
| Cytochrome P450, E-class, group I | 2.1E-19 |
| Cytochrome P450, conserved site | 4.6E-19 |
| Cytochrome P450 | 1.1E-18 |
| Monooxygenase | 3.0E-18 |
| Heme binding | 2.4E-15 |
| Iron ion binding | 1.0E-14 |

(B)
| GO term | P-value |
| --- | --- |
| GTP-binding | 2.1E-19 |
| P-loop containing nucleoside triphosphate hydrolase | 1.4E-8 |
| Nucleotide binding | 3.0E-8 |
| Guanylate-binding protein, C-terminal | 3.2E-6 |
| Guanylate-binding protein, N-terminal | 4.1E-6 |
| Septin | 7.5E-6 |

(C)
| GO term | P-value |
| --- | --- |
| Immunoglobulin C1-set | 1.4E-15 |
| MHC classes I/II-like antigen recognition protein | 2.2E-12 |
| Immunoglobulin/major histocompatibility complex, conserved site | 1.6E-9 |
| Immunoglobulin-like fold | 2.4E-9 |
| Immunity | 4.8E-9 |
| Allograft rejection | 6.9E-9 |

### 3.6. *M. sinica* only non-synonymous variants

We found 6614 autosomal heterozygous non-synonymous variant sites only detected in the *M. sinica* genome (*sinica*-specific). At these sites all other individuals used in the study were homozygous to the reference genome. Chromosome 1, 7, 14 and 19 and 20 showed higher levels of non-synonymous heterozygous sites compared to other chromosomes (Supplementary Table 1). In order to find if some chromosomes have more non-synonymous variants because of a biological impact or just because of the size of chromosome we determined the correlation between the length of the chromosomes and the number of variants found. We found that there is a slight positive correlation (Pearson correlation coefficient: r= 0.5635) between the number of sites and the length of the chromosome (according to reference genome chromosome length). Therefore more sites might not necessarily mean that a particular chromosome went through more mutations than others.

We conducted the GO (gene ontology) analysis for the genes that harbored variant sites and further looked at the top 10 genes that contained the most number of variants (Table 4). Of the top 10 genes with most variants 5 genes were associated with skeletal muscle formation and mutations of these were causing various diseases associated with muscle formation.

**Table 4.** Top 10 genes with the most number of non-synonymous SNPs found only in *M.sinica*. ‘ – ‘used where Gene function is not known or documented. The function or GO term of the genes are according to GeneCards.

| Gene | Expected function |
| --- | --- |
| WW domain binding protein 11 (WBP11) | Encodes for a protein that is associated with co-localizing with mRNA spicing factors and intermediate filament containing networks |
| Cardiomyopathy Associated 5 (CMYA5) | Anchor protein that mediates the subcellular compartmentation of protein kinase A (PKA)<br>Function as a repressor of calcineurin mediated transcription and negatively modulate skeletal muscle regeneration |
| Obscurin (OBSCN) | Organization of myofibrils during assembly<br>Mediate interactions between the sarcoplasmic reticulum and myofibrils |
| Zinc Finger Protein 721 (ZNF721) | Transcription regulation |
| ENSMMUG00000042203 | Uncharacterized protein |
| Solute carrier family 28 member 1 (CNT1) | Nucleoside binding and nucleoside:sodium symporter activity |
| Spectrin Beta (SPTBN5) | Actin and Kinesin binding |
| Zinc Finger Protein 469 (ZNF469) | Regulator for synthesis or organization of collagen fibers |
| Spectrin Repeat Containing Nuclear Envelope Protein 1 (SYNE1) | Forms a network between organelles and actin cytoskeleton |
| Stereocilin (STRC) | Associated with sensory hair cell, hair bundle in inner ear |

**Table 5.**
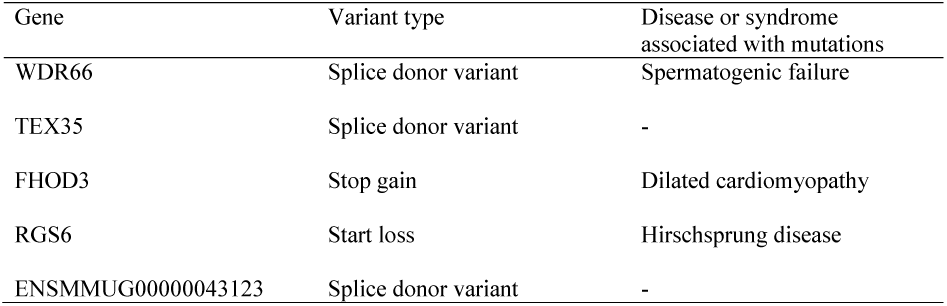
Five genes with high impact mutations in *M. sinica.* Mutations in some of these genes are associated with diseases. In the table ‘– ‘represents where there is no documented evidence of any disease or malfunction associated with mutation of the gene. Gene and disease information is from GeneCards.

In total, there were 20 HIGH impact *sinica*-specific non-synoymous mutations and out of those 5 were HIGH impact loss of function variant sites in 5 different genes (WDR66, TEX35, ENSMMUG00000043123 (uncharacterized), FHOD3 and RGS6) in *M. sinica* genome (Table 6). WDR66, TEX35, and ENSMMUG00000043123 had intron splice donor variants, while FHOD3 and RGS6 had stop codon gain and a start codon loss, respectively that were causing high impact on the corresponding genes. According to functional annotations and characterization regards to the human genome, WDR66 is a gene associated with sperm production and mutations in coding region cause spermatogenic failure, causing infertility. TEX35 is a testis expressed gene. The splice donor mutations normally lead to single exon skipping and defective protein formation (Abramowicz and Gos 2018) where the gene is unable to continue with its normal function. Defects in FHOD3 is associated with dilated cardiomyopathy (DCM), where the left ventricle of the heart becomes enlarged to the level that the heart fails to pump blood as normal (GeneCards). Stop codon gained mutation causes the protein to be short and may lead to loss of function of the protein. RGS6, which showed loss of start codon, is a gene responsible for regulation of G protein signaling and mutations cause Hirschsprung disease (HSCR) with large intestine or colon (Human Disease Database). Start loss mutations result in reduction or complete loss of function of the protein. Therefore there is a possibility that these genes may have been malfunctioning in this sequenced *sinica* individual.

## 4. Discussion

We sequenced the exome of *M. sinica*, an endemic macaque species to Sri Lanka for the first time. This attempt was to close the gap in the availability of sequence data of this species and also to understand its underlying genomic properties from a comparative genomics point of view. Here we found that *M. sinica* shared less polymorphisms with rest of the macaque species used in the study which included crab-eating and rhesus macaques. Similarly Tibetan macaque also showed a similar pattern with regards to polymorphisms. There were less heterozygous sites and also we found that the ratio of non-synonymous to synonymous SNPs were relatively higher in two *Sinica* species. Less genetic diversity is expected in isolated island populations such as *M. sinica*. The small restricted population size and isolation leads to inbreeding and then again reduced genetic variation. Tibetan macaques follow a similar pattern (Fan et al. 2014).

The phylogenetic tree construction and the PCA analysis both showed that *Mulatta* and *Fascicularis* cluster together forming one group, which is documented as *Fascicularis* group (Li et al. 2010) and *M. sinica* and *M. thibetana* clustered together in forming the *Sinica* group.

We found that the shared annotated high impact missense mutations of the 9 individuals used in the study reside in genes related to drug metabolism and MHC region. The non-synonymous sites found in *M. sinica* were found in genes related to zinc finger proteins. Also 5 of the the *M. sinica* only heterozygous non-synoymous sites revealed to have high impact loss of function effect on the genes they were located.

Here we presented the exome of *M. sinica* genome and also we looked at the exome sequences from numerous perspectives, but further analysis and studies on whole genome sequences of *M. sinica* genome may reveal a plethora of interesting facts of this rather less explored island population.

## 5. Conclusion

For the first time we sequenced *Macaca sinica* exome with 50x average coverage. This exome revealed less polymorphisms in the *M. sinica* genome similar to *M. thibetana* which is also an endemic macaque population. This could be a consequence of geographical isolation. The phylogenetic and clustering analyses revealed that *M. sinica* clusters with *M. thibetana and M. assamensi*s, whereas *M. fascicularis* and *M. mulatta* form their own distinct cluster. This finding agrees with the taxonomical grouping standard that had been introduced in the early 1970s’ by Fooden. Also our result shows congruence with Fan et al. 2018 phylogeny where taxonomically unresolved stump-tailed macaque showed close genetic relation to *sinica* group.

## Acknowledgments

We thank all those who provided numerous constructive comments and support to this work. We also thank curators, veterinarians and caretakers for collecting genetic samples. The study was approved as Collaborative Research of the Japan Monkey Centre (#2015006). This study was partially supported by MEXT KAKENHI to N.S and JSPS KAKENHI to T.H. (#16K18630). The toque macaque photograph was kindly provided by Associate Prof. Naoki Osada.

**The authors declare no conflict of interest.**

## Ethical approval

All applicable international, national, and/or institutional guidelines for the care and use of animals and genomic data were followed.

