## Supplementary Figures and Tables for "Exome sequences of toque macaques (*Macaca sinica*) of Sri Lanka reveal many amino acid changes"

Supplementary Material

Figure 1 – Total number of heterozygous, heterozygous synonymous and heterozygous non-synonymous sites found in the 9 individuals in the study.

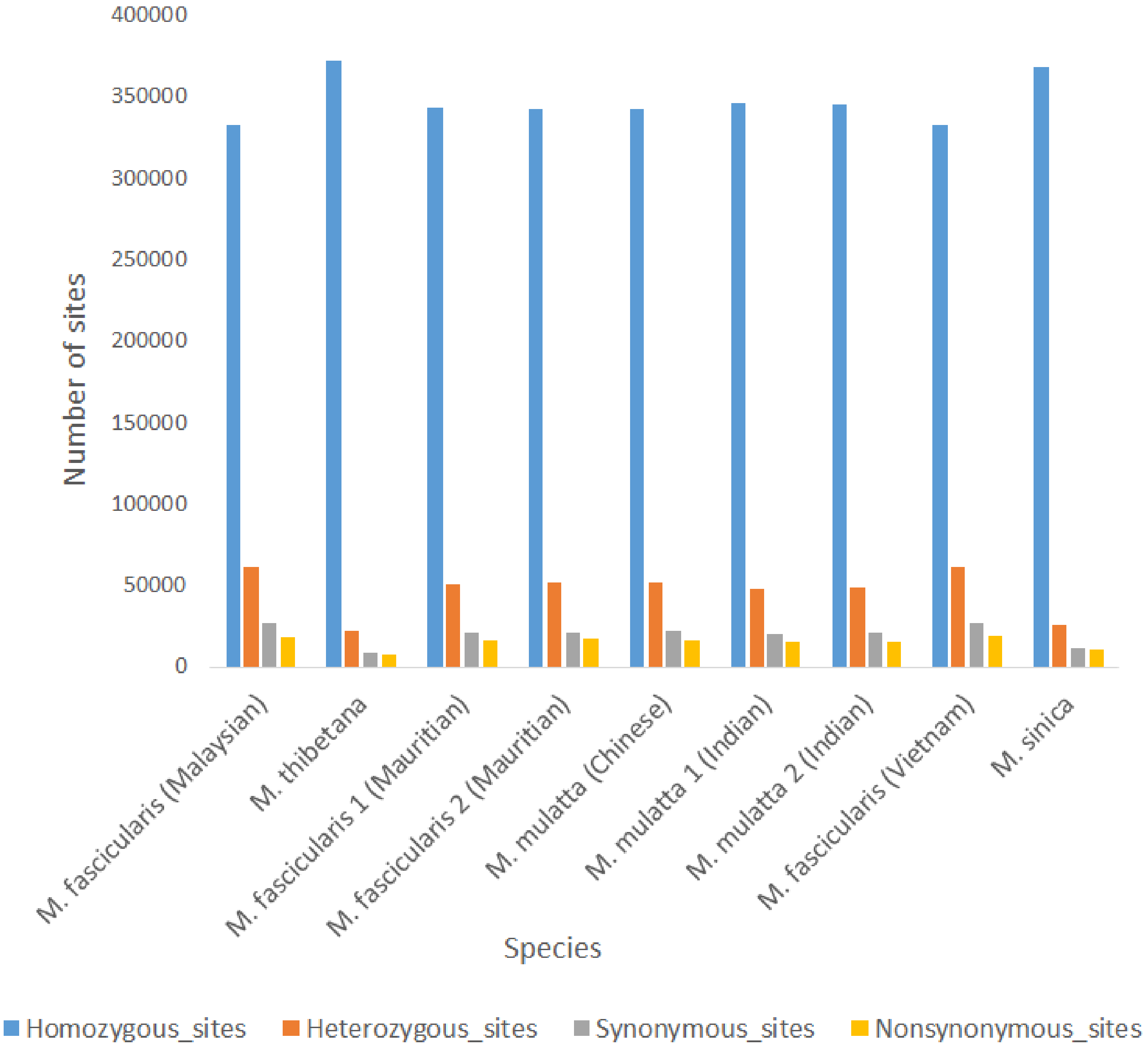

Figure 2 – Principal Component Analysis for 11 individuals after second variant calling.

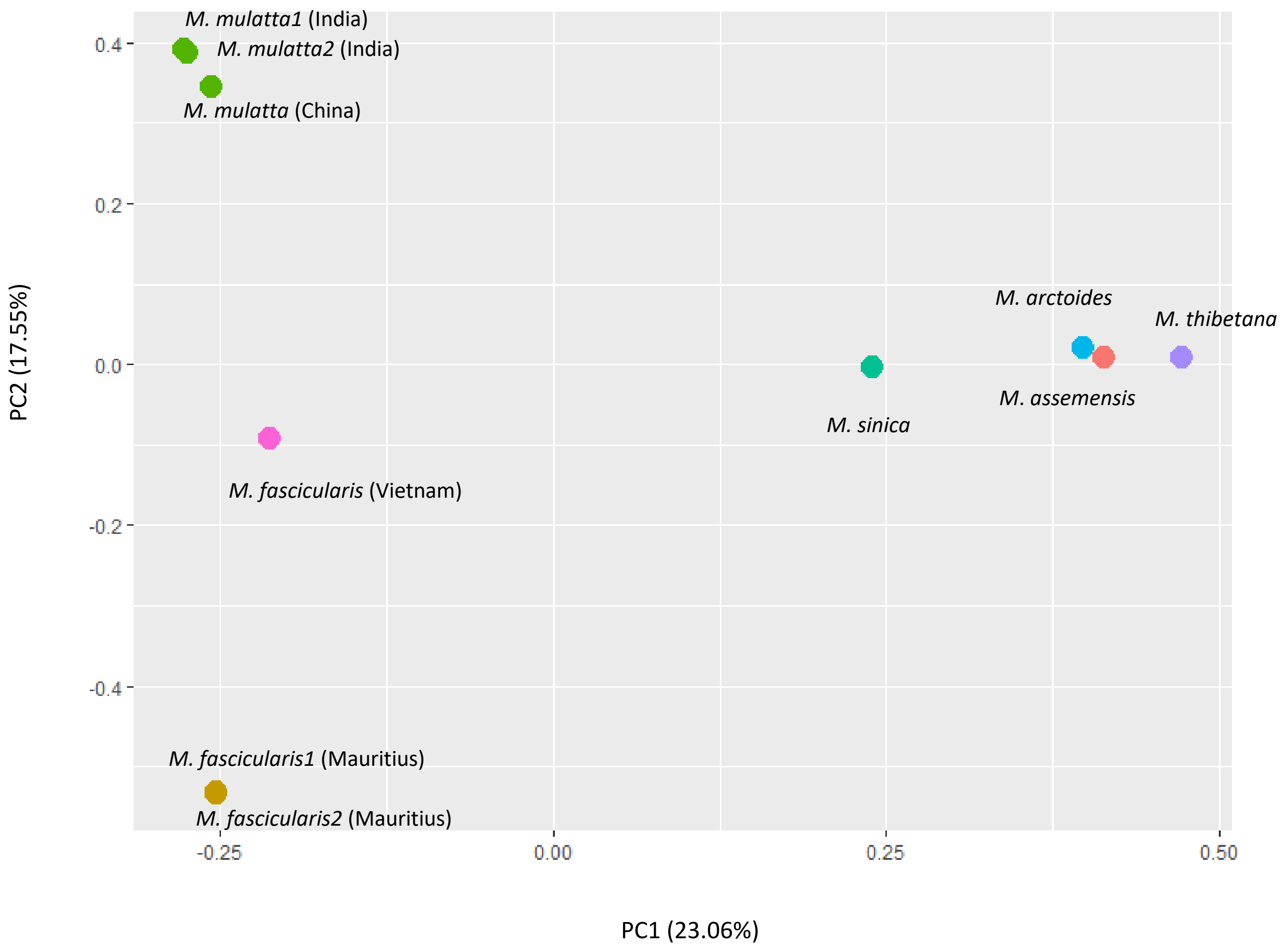

Table 1 – Heterozygosity levels of the 9 individuals after first variant calling

| Sample | Invariant sites | Heterozygous | Homozygous | Heterozygosity |
|---|---|---|---|---|
| *M. fascicularis* (Malaysia) | 56600355 | 113313 | 597840 | 0.001977 |
| *M. thibetana* | 58790767 | 41217 | 669925 | 0.00069 |
| *M. fascicularis* (Mauritius) | 58596012 | 95407 | 615699 | 0.001608 |
| *M. fascicularis* (Mauritius) | 58561338 | 98126 | 612872 | 0.001655 |
| *M. mulatta* (China) | 59258477 | 97667 | 613035 | 0.00162 |
| *M. mulatta* (India) | 59286265 | 91399 | 618617 | 0.001523 |
| *M. mulatta* (India) | 59179073 | 92682 | 615134 | 0.00154 |
| *M. fascicularis* (Vietnam) | 58891442 | 116120 | 585634 | 0.001948 |
| *M. sinica* | 32552736 | 116120 | 368076 | 0.00079 |

Table 2 – Number of heterozygous non-synonymous variant sites found only in *M. sinica* genome.

| Chromosome | Number of sites |
|---|---|
| 1 | 801 |
| 2 | 343 |
| 3 | 363 |
| 4 | 373 |
| 5 | 354 |
| 6 | 321 |
| 7 | 560 |
| 8 | 245 |
| 9 | 320 |
| 10 | 229 |
| 11 | 399 |
| 12 | 225 |
| 13 | 160 |
| 14 | 438 |
| 15 | 206 |
| 16 | 194 |
| 17 | 151 |
| 18 | 118 |
| 19 | 461 |
| 20 | 353 |
